# Synaptotagmin-9 and Tomosyn-1 molecular complex regulates Stx1A SNAREs to inhibit insulin secretion from pancreatic β-cells

**DOI:** 10.1101/2022.10.20.513128

**Authors:** Md Mostafizur Rahman, Asmita Pathak, Kathryn L Schueler, Haifa Al Sharif, Ava Michl, Justin Alexander, Jeonga-a Kim, Edwin Chapman, Sushant Bhatnagar

**Affiliations:** Heersink School of Medicine, Division of Endocrinology, Diabetes, & Metabolism, Comprehensive Diabetes Center, University of Alabama, Birmingham, AL, 35294; Department of Biochemistry, University of Wisconsin, Madison, WI, 53706; Investigator, Howard Hughes Medical Institute, Department of Neuroscience, University of Wisconsin, Madison, WI, 53706

**Keywords:** Synaptotagmin-9, Insulin Secretion, Tomosyn-1, SNARE, Syntaxin-1A, Exocytosis, Beta-cells, Islets, Insulin Granules, Biphasic Insulin Secretion

## Abstract

Stimulus-coupled insulin secretion from β-cells involves the fusion of insulin granules to the plasma membrane (PM) via SNARE complex formation—a cellular process key for maintaining whole-body glucose homeostasis. Optimal insulin secretion depends on how the clamping of SNAREs is released, rendering granules fusogenic. We show that an insulin granule protein synaptotagmin-9 (Syt9) deletion in lean mice increased glucose clearance, random-fed plasma insulin levels, and insulin secretion (in vivo and ex vivo islets) without affecting insulin sensitivity. These outcomes demonstrate that Syt9 has an inhibitory function in insulin secretion. Moreover, Syt9 interacts with PM-Stx1A and soluble Tomosyn-1 proteins to form non-fusogenic complexes between PM and insulin granules, preventing Stx1A-SNARE formation and insulin secretion. Furthermore, Syt9 inhibits SNARE-complex formation by posttranscriptional regulation of Tomosyn-1. We conclude that Syt9 and Tomosyn-1 are endogenous inhibitors that modulate Stx1A availability to determine β-cell secretory capacity.

**Highlights:**

- Synaptotagmin-9 inhibits biphasic insulin secretion from β-cells.
- Synaptotagmin-9, syntaxin-1A, and Tomosyn-1 forms a molecular complex that decreases the availability of syntaxin-1A to form SNARE complexes in insulin secretion.
- Synaptotagmin-9–mediated inhibition of insulin secretion occurs through post-transcriptional regulation of Tomosyn-1.

## Introduction

Glucose elicits a biphasic insulin secretion response from pancreatic β-cells to maintain wholebody glucose homeostasis^1^. An increased ATP to ADP ratio resulting from cellular glucose metabolism causes the closure of ATP-sensitive potassium channels (K_ATP_), leading to depolarization of the plasma membrane (PM)^2–4^. Consequently, this promotes the influx of Ca^2+^ by the L-type voltage-gated calcium channels, facilitating the fusion of insulin granules pre-docked or juxtaposed to the PM, releasing insulin primarily in the early phase^5^. Moreover, the fusion of insulin granules that traffic from inside of the cytoplasm contributes to the sustained phase^6^. Metabolic perturbations that reduce the ability of insulin granules to undergo PM fusion cause the development of β-cell dysfunction and Type 2 diabetes (T2D). However, the molecular mechanisms underlying the fusion of insulin granules in biphasic insulin secretion are not well characterized.

The soluble N-ethylmaleimide-sensitive factor attachment receptor protein (SNARE) complex formation is required for insulin granules to undergo fusion with the PM in both phases of glucose-stimulated insulin secretion (GSIS)^7–9^. Biochemical and structural studies have demonstrated that the SNARE core complex assembly occurs between a vesicle (v)-SNARE protein Vamp2 (vesicle-associated protein), which is present on the insulin granules with the PM-bound target (t)-SNARE proteins syntaxins (Stx) and Snap25 (synaptosome-associated protein of 25 kDa)^10^. Increases in [Ca^2+^]_i_ cause complete zippering of the SNAREs, accelerating membrane fusion and insulin exocytosis^11^. Stx1A is a well-characterized t-SNARE that forms a cognate SNARE core complex with Vamp2 and Snap25 to regulate the fusion of insulin granules in the early and sustained phases of insulin secretion^12–14^. Clusters of Stx1A are present in the PM with proximity to insulin granules^15^. Moreover, mice with β-cell-specific Stx1A deletion exhibit impaired early and sustained phase insulin secretion^12,16,17^. In the islets of individuals with T2D with reduced biphasic insulin secretion, the abundance of islet Stx1A is reduced^18^, implicating a clinically relevant role of Stx1A in β-cell function. Several proteins bind and modulate the function of Stx1A in forming SNARE complexes. Munc13-1^19^ and Munc18a^13^ facilitate the availability of Stx1A. In contrast, a relatively less characterized syntaxin binding protein 5 (Stxbp5) or Tomosyn-1 functions to clamp Stx1A, decreasing Stx1A-SNAREs (Stx1A-Snap25-Vamp2) formation and causing inhibition of insulin secretion^20^.

Synaptotagmin (Syt) isoforms have diverse roles in different cell types. The primary structure of Syt comprises an N-terminal intra-vesicular transmembrane region, a linker domain, and two Ca^2+^-binding cytoplasmic domains (C2A and C2B) at the C-terminal^21^. In the presence of Ca^2+^, Syt isoforms such as Syt1 and Syt7, via C2A and C2B, bind SNAREs and PM anionic phospholipids, accelerating SNARE complexes-mediated fusion of granules with the PM in exocytosis^22–25^. Additionally, Syt1 also functions to clamp SNARE fusion complexes^26^, and the ablation of Syt1 was found to increase the spontaneous release of granules from neurons in mice and drosophila^27–31^. In β-cells, Syt7 is a major Ca^2+^ sensor that facilitates insulin exocytosis^32–34^. Mice with the Syt7 deletion exhibited impaired insulin secretion and glucose intolerance^35^. However, the characterization of Syt isoforms that have roles in the clamping of SNARE complexes in β-cells remains elusive. The Syt9 (NCBI accession # NP_001347350, also referred to as Syt5^36^) is abundantly expressed in β-cells^37^. Also, it marks the sub-population of insulin granules with distinct release characteristics and lipid and protein compositions, compared to the Syt7 insulin granules^38^. Studies using clonal INS1 β-cells found that Syt9 increases insulin secretion^39,40^, whereas mice with pancreas-wide deletion of Syt9 showed no effect on insulin secretion and glucose clearance^41^. Thus, the role of Syt9 in insulin exocytosis remains undefined. Herein, we used the Syt9 gene deletion mouse model and clonal INS1(832/13) β-cells combined with metabolic phenotyping, confocal imaging, and biochemical approaches to elucidate the mechanism by which Syt9 regulates insulin secretion and whole-body glucose homeostasis.

## Results

### Loss of Syt9 improves glucose clearance associated with hyperinsulinemia

The deletion of Syt9 in *Syt9*^-/-^ vs. *Syt9*^+/+^ littermate control mice was confirmed by Western blotting. The protein abundance of Syt9 was undetectable in islets and brain of *Syt9*^-/-^ mice (Figure 1A). Metabolic phenotyping was performed using male *Syt9*^-/-^ and *Syt9*^+/+^ mice. No effect on body weight was observed regardless of the genotypes (Figure 1B). We then determined random-fed and fasting plasma insulin and glucose levels in *Syt9*^-/-^ and *Syt9*^+/+^ control mice on the standard chow diet. No significant difference in the random-fed (8 AM) blood glucose levels were observed (P = 0.06 and P = 0.33 at 6 and 10 weeks, respectively) (Figure 1C), whereas plasma insulin levels were increased by 2-fold (P = 0.001) (Figure 1D) at both 6 and 10 weeks in *Syt9*^-/-^ mice. No significant changes in 6 h fasting blood glucose levels were observed at 6 and 10 weeks in *Syt9*^-/-^ mice (Figure S1A). However, a small yet significant (1.3-fold, P = 0.01) increase in plasma insulin levels was observed at 6 weeks in *Syt9*^-/-^ mice, while no change was observed at 10 weeks. (Figure S1B). These data show that the primary effect of Syt9 loss is in increasing plasma insulin levels in the fed state. To evaluate the effect of Syt9 on insulin-stimulated glucose clearance, an oral glucose tolerance test was performed in *Syt9*^-/-^ and *Syt9*^+/+^ male mice. The loss of Syt9 led to accelerated glucose clearance (Figure 1E), reflected by the reduction in the glucose area under the curve (AUC) by 20% (P < 0.0001) that was observed in *Syt9*^-/-^, compared to the *Syt9*^+/+^ mice (Figure 1F). An insulin tolerance test was performed to evaluate the effect of Syt9 loss on insulin-stimulated glucose clearance in *Syt9*^-/-^ vs. *Syt9*^+/+^ control mice (Figure 1F). Neither the glucose excursions nor glucose AUCs for *Syt9*^-/-^ and *Syt9*^+/+^ mice after insulin injection were significantly different (Figures 1G and 1H, respectively), suggesting that the loss of Syt9 did not affect insulin action. Together, these data implicate the role of Syt9 in modulating pancreatic β-cell function. We also performed metabolic phenotyping in *Syt9^-/-^* and *Syt9*^+/+^ female mice. No pronounced effects on glucose homeostasis were observed; fasting and fed plasma insulin and glucose levels, glucose clearance, and insulin-stimulated glucose clearance (Figures S1C-S1I) remained unaffected by Syt9-deletion, suggesting a sex-specific role of Syt9 in regulating pancreatic β-cell function.

**Figure 1.**
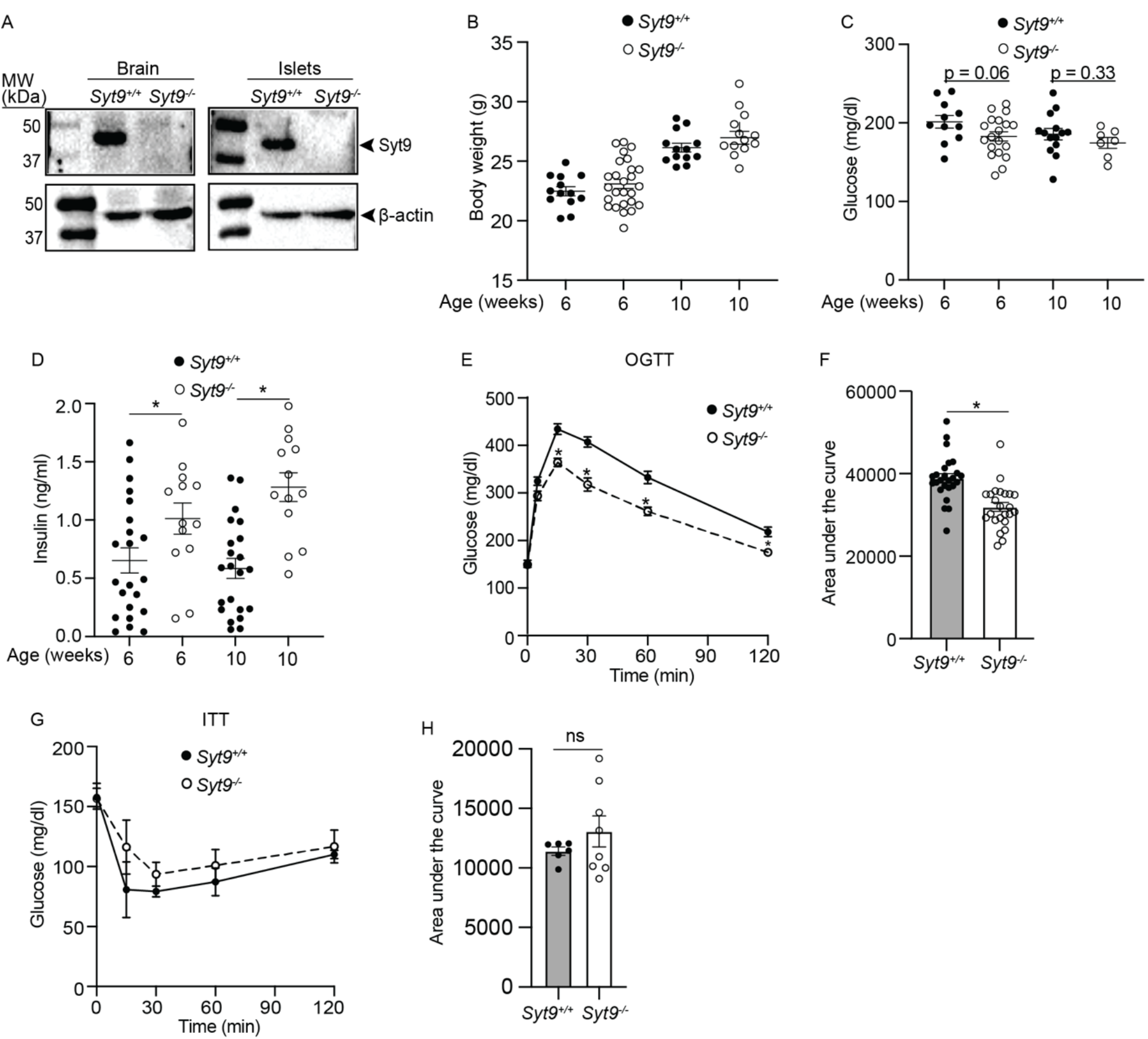
Mice lacking Syt9 shows improved glucose clearance. (A) Representative image of Western blots showing Syt9 and β-actin relative protein abundance in the brain (left) and islets (right) tissues from *Syt9*^-/-^ and *Syt9*^+/+^ mice (n > 4). (B) Body weight, (C) random-fed blood glucose levels, and (D) random-fed plasma insulin levels measurements were performed in *Syt9*^+/+^ and *Syt9*^-/-^ male mice at 6 and 10 weeks (n > 10). (E) Oral glucose tolerance test (OGTT) and (F) quantitation of the glucose AUCs encompassing 120 min in GTT was performed in *Syt9*^+/+^ and *Syt9*^-/-^ male mice (n > 20). G) Insulin tolerance test (ITT) and (H) quantitation of the glucose AUCs in ITT was performed in *Syt9*^+/+^ and *Syt9*^-/-^ male mice (n > 6). Data are presented as mean ± SEMs. *P < 0.05; **P < 0.01; ***P < 0.001

### Loss of Syt9 increases stimulus-coupled insulin secretion

Increases in plasma insulin levels were observed in random-fed *Syt9*^-/-^ mice. Thus, we investigated the effect of Syt9 loss on insulin secretion. The *Syt9*^-/-^ and *Syt9*^+/+^ mice were subjected to an oral glucose challenge, and plasma insulin levels were determined at different time points (Figure 2A). Glucose increased plasma insulin levels at 5 and 15 min compared to the baseline (t = 0 min) in both groups; however, further increases by ~30% (P = 0.04) and ~65% (P = 0.03) were observed in *Syt9*^-/-^ mice compared to *Syt9*^+/+^ mice (Figure 2A). These data suggest that increases in insulin secretion contribute to improved glucose clearance in Syt9-deficient mice.

**Figure 2.**
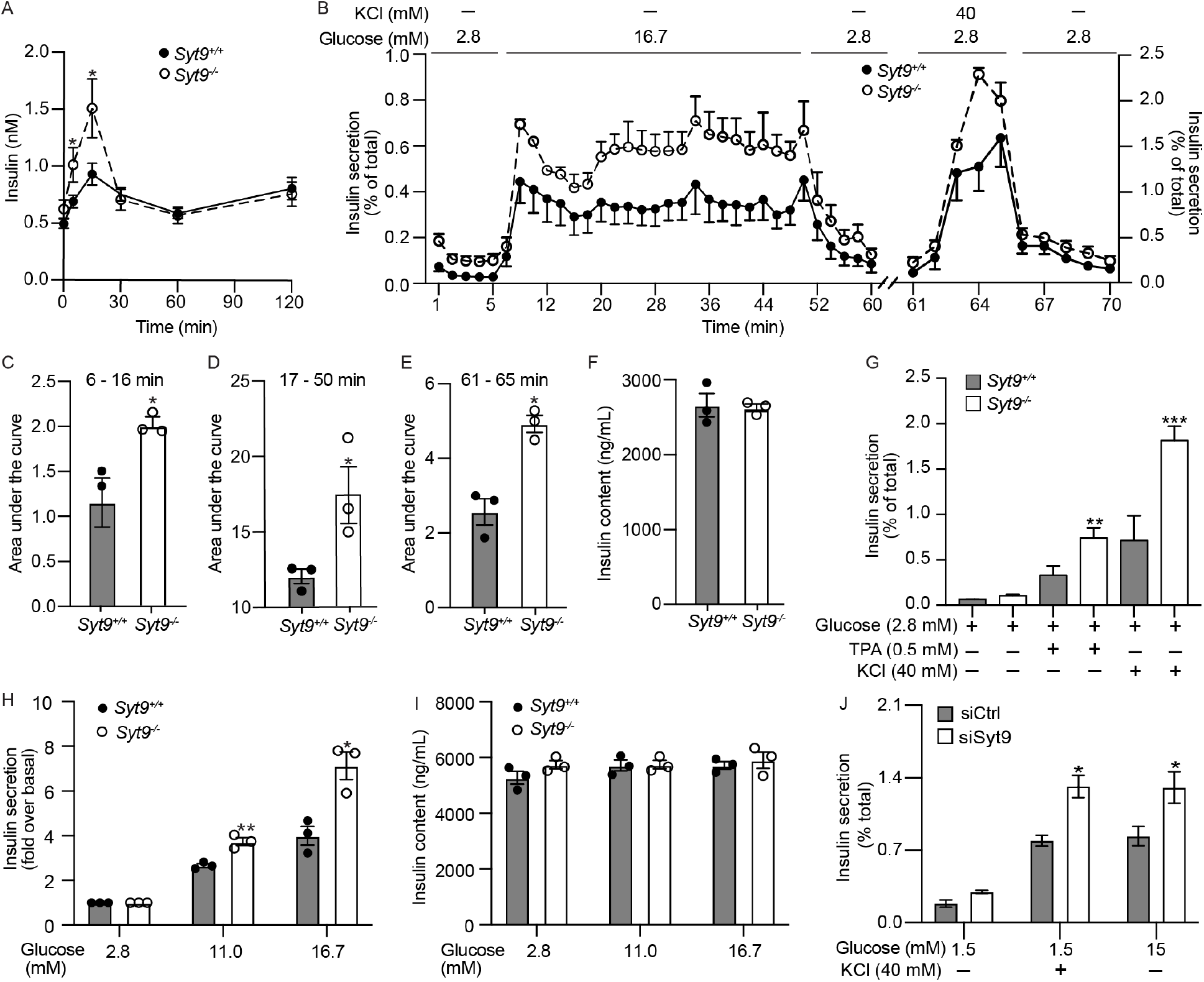
The loss of Syt9 increases biphasic insulin secretion. (A) Plasma insulin levels during OGTT in *Syt9*^+/+^ and *Syt9*^-/-^ male mice (n > 20). (B) Perifusion to determine dynamic insulin secretion from islets isolated from *Syt9*^+/+^ and *Syt9*^-/-^ mice (N = 3). Insulin secretion AUC for (C) early and (D) sustained phase in response to glucose. (E) Insulin secretion AUC in response to KCl. (F) Insulin content of islets subjected to perifusion. (G) Static insulin secretion from islets isolated from *Syt9*^+/+^ and *Syt9*^-/-^ mice (n = 4) in response to the stimulation by KCl and TPA (12-O-tetradecanoylphorbol-13-acetate). (H) Static insulin secretion and (I) insulin content from *Syt9*^+/+^ and *Syt9*^-/-^ mouse islets (n = 4) in response to glucose dose (2.8, 11, 16.7 mM). (J) Static insulin secretion from INS1(832/13) clonal β-cells transfected with siScramble or siSyt9 in response to stimulation by glucose or KCl for 15 min (n = 3). Data are presented as mean ± SEMs. *P < 0.05; **P < 0.01; ***P < 0.001

To gain insights into how Syt9 regulates insulin secretion, we performed dynamic insulin secretion measurements by perifusion^42^ using islets isolated from *Syt9*^-/-^ and *Syt9*^+/+^ mice (Figures 2B–2I). As expected, high glucose (16.7 mM) elicited increases in early (6 – 16 min) and sustained (17 – 50 min) phases of insulin secretion compared to low glucose (2.8 mM) in both groups of mouse islets (Figure 2B). However, the *Syt9*^-/-^ mouse islets exhibited higher increases in early and sustained phase insulin secretion compared to *Syt9*^+/+^ mouse islets (Figure 2B). These are reflected by the AUCs, which revealed a two-fold increase in the early (Figure 2C, P = 0.04) and a 50% increase in the sustained phase (Figure 2E, P = 0.04) insulin secretion from *Syt9*^-/-^ vs. Syt*9*^+/+^ mouse islets in response to high glucose. KCl is known to cause PM depolarization and facilitates insulin release in the early phase. Consistently, KCl (40 mM)-induced early-phase insulin secretion was observed from both *Syt9*^-/-^ vs. *Syt9*^+/+^ mouse islets (Figure 2B). However, islets from *Syt9*^-/-^ mice exhibited significantly increased insulin secretion compared to *Syt9*^+/+^ mouse islets (Figure 2B), as reflected by the 2-fold increase in the AUC for KCl-stimulated insulin secretion profile (Figure 2E, P = 0.004). These outcomes show that Syt9 functions as an inhibitor of biphasic insulin secretion.

Static insulin secretion was performed using islets isolated from *Syt9*^-/-^ vs. *Syt9*^+/+^ mice to gain insights into Syt9-regulated insulin secretion. The 0.5 mM phorbol ester (TPA) (2-fold, P = 0.001) and 40 mM KCl (3-fold, P = 0.0007) at basal concentrations of glucose (2.8 mM) stimulated insulin secretion was significantly increased in *Syt9*^-/-^ mouse islets compared to the *Syt9*^+/+^ mouse islets.

Similar to islet perfusion outcomes, significant increases in the 11 mM glucose (1.5-fold, P = 0.005) and 16.7 mM glucose (1.8-fold, P < 0.01) stimulated insulin secretion were observed in *Syt9*^-/-^ vs. *Syt9*^+/+^ mouse islets (Figures 2G–2H). Moreover, Syt9 deletion did not affect basal insulin secretion at 2.8 mM glucose (Figures 2B, 2F, 2G, 2H), suggesting that Syt9 regulates stimulus-coupled insulin secretion. Also, islet insulin content remained unchanged upon Syt9 deletion (Figures 2F, 2I), implicating that Syt9 inhibits insulin secretion by modulating the insulin secretory pathway or exocytosis. Furthermore, Syt9 deletion increased insulin secretion in response to metabolizable (glucose) and non-metabolizable (TPA and KCl) insulin secretagogues, indicating that Syt9 functions by regulating distal steps in the insulin exocytosis.

To confirm the role of Syt9 in insulin secretion, a small inhibitory (si) RNA approach was used to achieve the knockdown of Syt9 in clonal INS1(832/13) β-cells. Greater than 90% reduction in Syt9 mRNA (Figure S2A) and protein levels (Figure 4I, Syt9 input blot) were achieved upon Syt9 knockdown in INS1(832/13) cells. As in islets, basal insulin secretion and β-cell insulin content were unaffected (Figures S2B-S2C). However, increases in insulin secretion in response to the stimulation by KCl (40 mM at 1.5 mM glucose) and high glucose (15 mM) were observed in Syt9-knockdown compared to siScramble INS1(832/13) cells (Figure 2J). Collectively, these outcomes demonstrate that Syt9 inhibits biphasic insulin secretion from β-cells.

**Figure 3.**
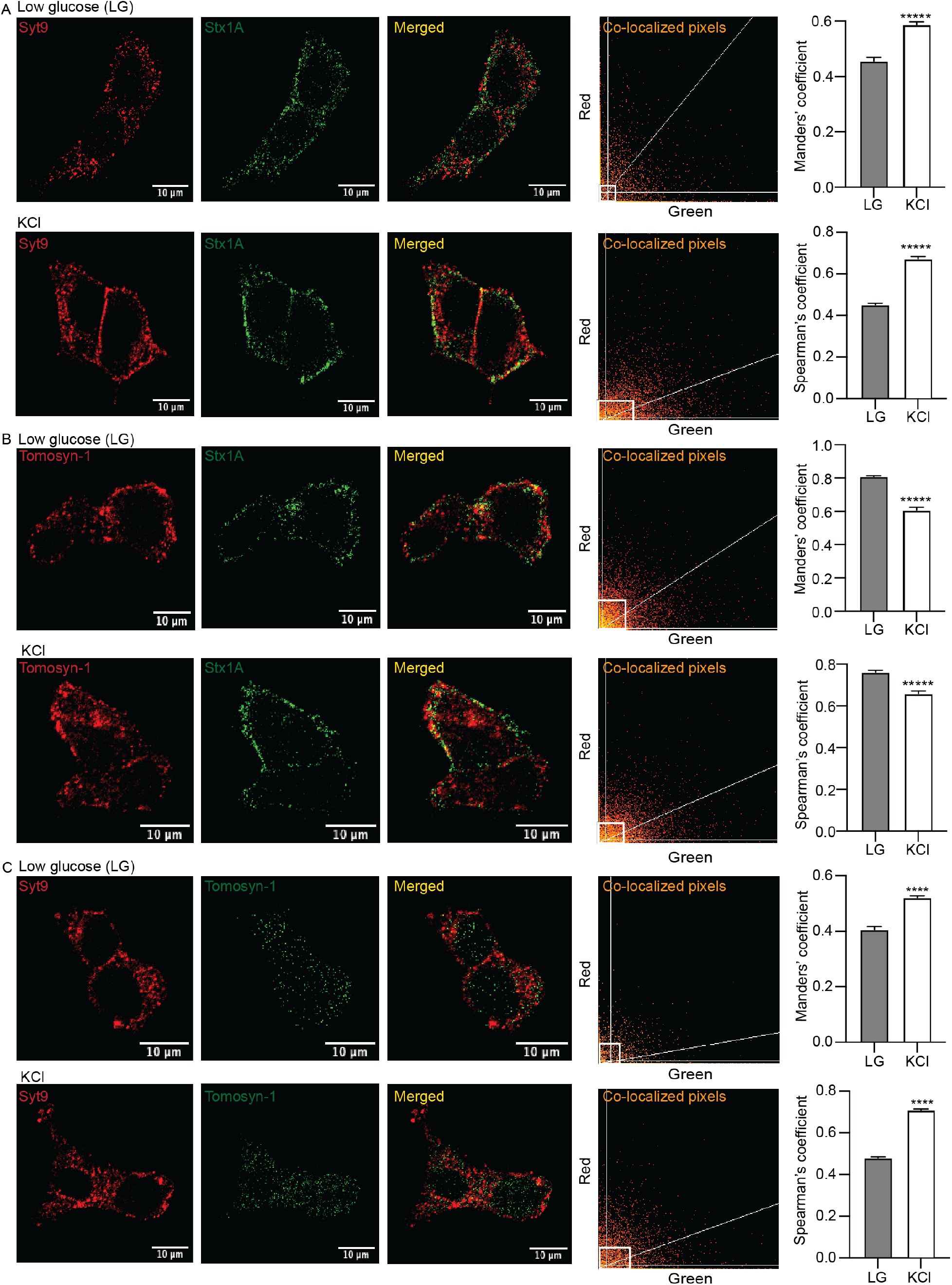
Colocalization of Syt9, Tomosyn-1, and Stx1A in pancreatic β-cell. Colocalization of (A) Syt9 (red) and Stx1A (green), (B) Tomosyn-1 (red) and Stx1A (green) and (C) Syt9 (red) and Tomosyn-1 (green) in INS1 (832/13) cells in response to low glucose (1.5 mM) or KCl (40 mM) treatment for 15 min was observed using immunofluorescence. Images were captured using confocal (scale bar = 10 μm) at 60X, and co-localized pixels were analyzed using Image J and estimated by Manders’ and Spearman’s coefficients. Data are presented as mean ± SEMs. *P < 0.05; **P < 0.01; ***P < 0.001

**Figure 4.**
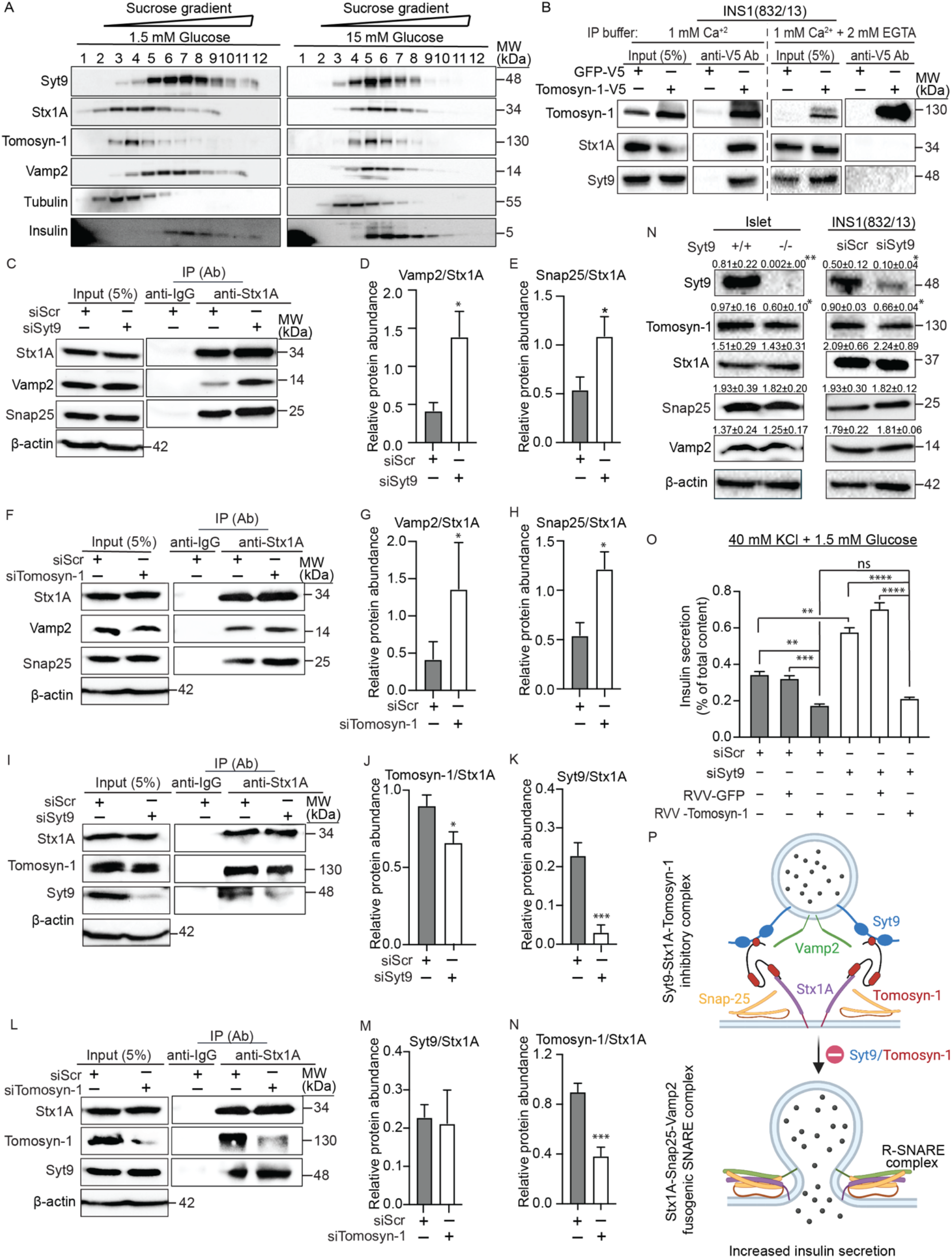
Syt9 forms a tripartite complex with Tomosyn-1 and Stx1A to inhibit SNARE complex formation. (A) Cofractionation of INS1 (832/13) cells by sucrose gradients prepared at low (1.5 mM) and high (15 mM) glucose (n = 3). (B) Immunoprecipitation (IP) was performed in INS1(832/13) cells using an anti-V5 antibody. IP was performed in buffer supplemented with 1 mM Ca^2+^ (Left) or 2 mM EGTA to chelate Ca^2+^ (right) (n = 3). (C) Co-IP by anti-Stx1A antibody using lysates prepared from INS1(832/13) cells transfected with siScramble (Scr) and siSyt9 after stimulation by KCl for 15 min. (D) Relative binding of Vamp2 protein to Stx1A (n=5). (E) Relative binding of Snap25 (right) to Stx1A (n = 5). (F) Co-IP by anti-Stx1A antibody using lysates prepared from INS1(832/13) cells transfected with siScr and siTomosyn-1 after stimulation by KCl for 10 min. (G) Relative binding of Vamp2 protein to Stx1A (n=5). (H) Relative binding of Snap25 (right) to Stx1A (n = 5). (I) Co-IP by anti-Stx1A antibody using lysates prepared from INS1(832/13) cells transfected with siScr and siSyt9 after stimulation by KCl for 15 min. (J) Relative binding of Tomosyn-1to Stx1A (n=5). (K) Relative binding of Syt9 to Stx1A (n = 5). (L) Co-IP by anti-Stx1A antibody using lysates prepared from INS1(832/13) cells transfected with siScr and siTomosyn-1 after stimulation by KCl for 15 min. (M) Relative binding of Syt9 to Stx1A (n=5). (N) Relative binding of Tomosyn-1 to Stx1A (n = 5). (O) Protein abundance was determined in *Syt9*^+/+^ and *Syt9*^-/-^ mouse islets (left) and INS1 (832/13) cells (right). The number presented on the top of each band represents the protein abundance compared to β-actin (n > 4). (P) Insulin secretion (% of total) in response to KCl (40 mM) stimulation in INS1(832/13) cells overexpressing Tomosyn-1 with siScr or siSyt9 (n = 3). (Q) Top, Syt9 and Tomosyn-1 clamp Stx1A limiting the formation of the Stx1A-Snap25-Vamp2 SNARE complexes. Bottom, removing Tomosyn-1 and Syt9 facilitates Stx1A-Snap25-Vamp2 SNARE complexes in insulin secretion. Data are presented as mean ± SEMs. *P < 0.05; **P < 0.01; ***P < 0.001

### Syt9 colocalization increases with Stx1A and Tomosyn-1 upon stimulation

Stx1A forms SNARE complexes (Stx1A-Snap25-Vamp2) in insulin exocytosis^43^. Alternatively, Tomosyn-1 is an inhibitor of insulin exocytosis that clamps Stx1A, thereby limiting the formation of Stx1A-SNARE complexes^44,45^. Thus, we used confocal imaging to evaluate whether endogenous Syt9 colocalizes with Stx1A and Tomosyn-1 to regulate insulin secretion. Colocalization between proteins in INS1(832/13) cells after 15 min of treatment with low glucose (1.5 mM) and KCl (40 mM at 1.5 mM glucose) was estimated by Manders’ overlap coefficient (MOC) and Spearman’s coefficient (r). The staining of Syt9 (red) with Stx1A (green) is shown as a merge (yellow) (Figure 3A). Colocalization estimated by MOC and r between Syt9 and Stx1A was observed at low glucose and KCl stimulation. However, significant increases in the colocalization between Syt9 and Stx1A were observed upon KCl stimulation compared to low glucose (MOC_KCl_ = 0.6 vs. MOC_LG_ = 0.45, P < 0.0001; r_KCl_ = 0.65 vs. rLG, P < 0.0001) (Figure 3A). Similarly, we assessed the colocalization between Tomosyn-1 (red) and Stx1A (green) (Figure 3B). Colocalization shown as merge (yellow) was observed in response to low glucose and KCl stimulation. However, a slight but significant reduction in the colocalization between Tomosyn-1 and Stx1A was observed upon KCl stimulation compared to low glucose (MOC_KCL_ = 0.6 vs. MOC_LG_ = 0.8, P < 0.0001; r_KCL_=0.65 vs. r_LG_=0.78; P < 0.0001). These outcomes show an inverse relationship between the colocalization of Tomosyn-1 and Syt9 for Stx1A in β-cells upon stimulation. Next, we assessed the colocalization between Tomosyn-1 (green) and Syt9 (red) (Figure 3C). Colocalization between Tomosyn-1 and Syt9, shown as yellow/orange, was observed in response to low glucose and KCl stimulation. Surprisingly, significant increases between Syt9 and Tomosyn-1 colocalization were observed in response to KCl stimulation compared to low glucose (MOC_KCL_ = 0.5 vs. MOC_LG_ = 0.4, P < 0.001; r_KCL_ = 0.7 vs. r_LG_ = 0.45, P < 0.001). Altogether, the colocalization observed between Tomosyn-1, Stx1A, and Syt9 at low glucose culminated in increased colocalization between Syt9-Stx1A and Syt9-Tomosyn-1 upon stimulation. Upon stimulation, proportional increases in colocalization between Syt9-Stx1A and Syt9-Tomosyn-1 suggest that Tomosyn-1, Stx1A, and Syt9 form a molecular complex in β-cells.

### Loss of Syt9 increases the formation of Stx1A-SNARE complexes

We investigated the mechanism by which Syt9 regulates insulin secretion. Subcellular fraction of INS1(832/13) cells were prepared at low glucose (1.5 mM) and high glucose (15 mM) by sucrose density gradient centrifugation. Cofractionation was observed between Stx1A with Tomosyn-1 and of Syt9 with granule proteins Vamp2 and insulin at low glucose. Upon stimulation, Syt9, Stx1A, Tomosyn-1, Vamp2, and insulin were primarily present in the same fractions (Figure 4A). This outcome is consistent with the confocal imaging showing the colocalization between Syt9, Stx1A, and Tomosyn-1 upon stimulation. Next, we determined whether Syt9, Stx1A, and Tomosyn-1 form tripartite protein complexes. The INS1(32/13) cell lysates expressing V5-Tomosyn-1 were prepared after 15 min of stimulation by KCl (40 mM at 1.5 mM glucose). Overexpression caused a 4-fold increase in Tomosyn-1 protein abundance compared to endogenous levels (Figure 4B). Immunoprecipitation (IP) of Tomosyn-1 was performed using an anti-V5 antibody in IP buffer with 1 mM calcium, mimicking conditions conducive to forming SNARE complexes. The immunoprecipitated protein complex subjected to Western blotting showed that Tomosyn-1 binds specifically to Stx1A and Syt9 (Figure 4B). No bands for Tomosyn-1, Stx1A, and Syt9 were observed in protein complex immunoprecipitated by anti-V5 antibody using control INS1(832/13) cell lysates. Previously it was reported that the binding of Tomosyn-1 with Syt1 and Stx1A increased in the presence of calcium^46^. Conversely, chelating calcium with 2 mM EGTA led to Tomosyn-1 IP, but no binding was observed with Stx1A and Syt9. These data establish that Tomosyn-1 binds to Stx1A and Syt9 in β-cells upon stimulation.

To assess the inhibitory role of Stx1A-Tomosyn-1-Syt9 protein complex in insulin secretion, we determined whether the knockdown of Syt9 or Tomosyn-1 would increase formation of Stx1A-SNARE complexes (Snap25-Vamp2 SNARE) —required for the fusion of insulin granules to the PM in insulin exocytosis. INS1(832/13) cells, transfected with siSyt9, siTomosyn-1, and siScramble, were treated with KCl for 15 min before preparation of cell lysates. The protein abundance of Syt9 and Tomosyn-1 was reduced by 90% (Figure 4I, input blot) and ~75% (Figure 4L, input blot), respectively. Anti-Stx1A antibody was used to co-immunoprecipitate (Co-IP) Stx1A protein complex for Western blotting. Binding between Stx1A, Vamp2, and Snap25 was observed in siScramble cell lysates, confirming Stx1A-SNARE complex formation (Figure 4C). However, increases in the binding of Vamp2 and Snap25 with Stx1A were observed in siSyt9 or siTomosyn-1 cell lysates compared to siScramble control cell lysates (Figures 4C, 4F). Quantification showed significant enrichment of Vamp2/Stx1A (siSyt9/siScramble = 3-fold, P = 0.02; siTomosyn-1/siScramble = 3-fold, P = 0.01) and Snap25/Stx1A (siSyt9/siScramble = 2-fold, P = 0.04; siTomosyn-1 siTomosyn-1/siScramble = 2-fold, P = 0.01) upon knockdown of Syt9 (Figures 4D–4E) or Tomosyn-1 (Figures 4G–4H). The absence of Stx1A, Vamp2, and Snap25 in IP performed using isotype control IgG antibody showed specificity in IP Stx1A-SNARE proteins using anti-Stx1A antibody. These outcomes suggest that Syt9 and Tomosyn-1 bind and block Stx1A’s ability to form Stx1A-SNARE complexes for the PM fusion of insulin granules in exocytosis.

### Inhibitory effects of Syt9 on insulin secretion are mediated by Tomosyn-1

We evaluated the interrelationship between Syt9 and Tomosyn-1 in modulating Stx1A-SNARE complexes. INS1(832/13) cells transfected with siSyt9, siTomosyn-1, and siScramble for 42 h were subjected to 15 min of KCl stimulation before the preparation of cell lysates. Subsequently, immunoprecipitation with anti-Stx1A antibody was performed to assess the binding of Tomosyn-1 and Syt9 with Stx1A in response to Tomosyn-1 or Syt9 knockdown. The binding of Stx1A with Tomosyn-1 and Syt9 was observed in siScramble cell lysates (Figures 4I, 4L), confirming the formation of endogenous tripartite Stx1A, Tomosyn-1, and Syt9 protein complexes in β-cells. Interestingly, significantly reduced binding of Tomosyn-1 with Stx1A was observed upon Syt9 knockdown (1.4-fold decrease, P = 0.04) (Figures 4I, 4J). However, the binding between Stx1A and Syt9 was not affected by the Tomosyn-1 knockdown (Figures 4L, 4M). As expected, significantly reduced binding between Stx1A and Syt9 upon Syt9 knockdown (7-fold, P = 0.0009) (Figure 4K) and Stx1A and Tomosyn-1 upon Tomosyn-1 knockdown (2.4-fold, P = 0.0008) (Figure 4N) were observed due to reduction of Syt9 and Tomosyn-1 proteins, respectively. The IP performed using isotype control IgG antibody showed specificity in immunoprecipitating Stx1A-SNARE proteins. These outcomes suggest that Syt9 facilitates the binding of Tomosyn-1 to Stx1A, modulating the availability of Stx1A to form fusogenic Stx1A SNARE complexes.

Alterations in Tomosyn-1 protein abundance directly affect exocytosis^47^. Thus, we evaluated whether Syt9 loss/knockdown reduced Tomosyn-1 protein abundance. Cell lysates were prepared from a) islets and brain isolated from *Syt9*^-/-^ and *Syt9*^+/+^ mice and b) siScramble and siSyt9 transfected INS1(832/13) cells. A forty percent (P = 0.01) decrease in Tomosyn-1 protein abundance (Figure 4N) was observed without alterations in mRNA levels (Figure S3A-S3B) upon Syt9 loss/knockdown. Additionally, Stx1A, Snap25, or Vamp2 protein levels were not affected by Syt9 loss (Figure 4O). These outcomes suggest that Syt9 regulates Tomosyn-1 protein abundance post-transcriptionally. Also, reductions in Tomosyn-1 protein levels were observed in *Syt9^-/-^* brain tissue (Figure S3C), implicating that the role of Syt9 in regulating Tomosyn-1 protein is not due to β-cell specific signaling is potentially a consequence of Syt9–Tomosyn-1 interaction.

We assessed whether Syt9 effects on insulin secretion are mediated by Tomosyn-1. Insulin secretion was performed by overexpressing Tomosyn-1 in siSyt9 or siScramble INS1(832/13) cells treated with low glucose (1.5 mM), high glucose (15 mM), and KCl (40 mM at 1.5 mM glucose) for 15 min. KCl (Figure 4Q) and high glucose (Figure S4B) treatments increased insulin secretion compared to low glucose (Figure S4A) (9-fold increase, P = 0.0001). Tomosyn-1 inhibited KCl (P = 0.0001) and high glucose (P = 0.0001) stimulated insulin secretion by ~40%. The data presented in Figure 2J show that the knockdown of Syt9 increased insulin secretion in response to KCl and glucose by ~2-fold (5^th^ vs. 2^nd^ bar in Figures 4O, S4B). Interestingly, no increase in the KCl- or high glucose-stimulated insulin secretion was observed upon Syt9 knockdown with the overexpression of Tomosyn-1 (6^th^ vs. 3^rd^ bar in Figures 4O, S4B). The basal insulin secretion and cellular insulin content were not altered (Figures S4A, S4C). These outcomes demonstrate that Tomosyn-1 mediates increases in insulin secretion observed upon the loss or knockdown of Syt9.

## Discussion

Stx1A is a well-characterized t-SNARE required for SNARE complex-mediated fusion of insulin granules in the early and sustained phases of insulin secretion. Less is known about how β-cells clamp or inhibit Stx1A, limiting the formation of SNARE complexes in exocytosis. The outcomes of this study demonstrate the undescribed role of Syt9 as an endogenous inhibitor of biphasic insulin secretion that functions via Tomosyn-1 to regulate the formation of the Stx1A-SNARE complex (Figure 4P). Ablation of Syt9 in lean mice increased glucose clearance, random-fed plasma insulin levels, and insulin secretion (in vivo and ex vivo islets) without affecting insulin sensitivity (Figures 1–2). Syt9 knockdown in clonal β-cells also increased stimulus-coupled insulin secretion (Figures 2J, 4Q). Collectively, these results show that β-cell Syt9 directly regulates insulin secretion and glucose clearance. Elucidating the mechanism by which endogenous inhibitors modulate insulin secretion will provide insights into several unresolved facets of insulin exocytosis, such as: why only a fraction of cellular insulin granules undergo PM fusion upon stimulation^48^, aged-insulin containing granules are not preferred for fusion^49–52^, there is a loss of early-phase insulin secretion in impaired glucose tolerance^52–56^, and an increase in fusion-incompetent docked granules and reduced fusion competency of granules in T2D^9,10,57–63^.

Insulin granules from different cellular pools contribute to biphasic GSIS^64^. Pre-docked granules are juxtaposed to the PM, they immediately undergo fusion upon stimulation, and account for fifty percent of insulin released in the early phase^65^. In comparison, newcomer granules are present in the cytoplasm away from the PM^66^. Upon stimulation, they undergo fusion by two plausible modes, short-dock or no-dock, contributing to insulin released in both phases. Our results show that Syt9 deletion increased early and sustained phases of GSIS from islets (Figures 2B, 2F, 2G), implicating that Syt9 has a role in regulating the fusion of pre-docked and newcomer insulin granules.

Docking is a temporal constraint on granules^67^, and mechanisms underlying long and short duration docking of pre-docked and newcomer granules, respectively, are not completely understood. Newcomer granules are trafficked from within the cells to the PM. Thus, a potential role of an inhibitor that functions at the insulin granule–PM interface has been proposed. Stx1A is required for biphasic insulin secretion and facilitates the fusion of pre-docked and newcomer with short-dock insulin granules^12^. We show that Syt9 (an insulin granule protein) colocalizes, binds, and cofractionates with the PM Stx1A (Figures 3, 4A, 4I). These outcomes, combined with Syt9’s inhibitory function on insulin secretion (Figure 2), suggest that Syt9 potentially functions as a clamp of Stx1A, modulating its availability to form SNARE complexes-mediated fusion of predocked and newcomer (short-dock) insulin granules. How Syt9 modulates fusion modes of insulin granules involving distinct cognate SNARE complexes are part of the future directions.

It is proposed that Syt clamps SNARE complexes mediated fusion by either imposing a separation between the PM and granule membrane that is greater for the SNARE-complex-mediated fusion^68^ or by locking the SNAREs in a fusion incompetent state^69–71^. Insights into Syt functioning are predominately derived from studies elucidating the role of Syt1 in neurotransmitter release. However, less is known about Syt isoforms in regulating SNARE-mediated insulin exocytosis. Our data provide a mechanism by which Syt9 via Tomosyn-1 inhibits Stx1A-SNARE complexes in insulin exocytosis. Tomosyn-1 is a soluble cytoplasmic protein that inhibits SNARE complex formation^72^. It interacts with Stx1A via a short C-terminal domain, clamping Stx1A and Snap25 in a non-fusogenic complex. This leads to blocking the stimulatory effects of Munc13-1 and Munc18-1 in facilitating Stx1A-SNARE assembly^73,74^. Tomosyn-1 also associates with insulin granules via N-terminal domains^75–79^. Furthermore, deletion of the two unstructured loops in the N-terminal was found to block the inhibitory function of Tomosyn-1 while retaining Stx1A binding^45^, demonstrating that N-terminal domains are required for the complete inhibitory function of Tomosyn-1. Our data have identified that Syt9 (a granule protein), Tomosyn-1, and Stx1A (a PM protein) molecular complex is potentially formed at the PM–insulin granules interface (Figure 4B). As demonstrated for the Tomosyn-1-Syt1 binding^46^, the N-terminal domains in Tomosyn-1 may be responsible for binding to Syt9 in β-cells. Also, the knockdown of Syt9 or Tomosyn-1 increased the formation of Stx1A-Snap25-Vamp2 SNAREs (Figures 4C, 4F), establishing that Syt9 and Tomosyn-1 are inhibitors of Stx1A-SNARE complexes in insulin exocytosis. Thus, the Syt9-Tomosyn-1-Stx1A complex functions by decreasing the availability of Stx1A to form SNAREs, rendering insulin granules transiently or completely non-fusogenic.

The availability of Stx1A to form SNARE complexes increases insulin secretion. Our data provide a mechanism by which Syt9 inhibits insulin secretion. Syt9 ablation decreased Tomosyn-1 protein (not mRNA) levels and the binding of Tomosyn-1 with Stx1A (Figures 4N, 4I). Moreover, increases in Stx1A-Snap25-Vamp2 SNARE complexes were observed upon Tomosyn-1 knockdown (Figures 4F–4H); alterations in Tomosyn-1 protein abundance are known to affect Stx(s) function proportionally. Furthermore, Tomosyn-1 overexpression blocked increases in insulin secretion due to Syt9 knockdown from β-cells (Figure 4Q). We show that the loss of Syt9 by post-transcriptional mechanisms decreases Tomosyn-1 inhibitory function, thereby increasing Stx1A-SNARE-mediated insulin secretion. Post-translational regulation by Hrd1 E3-polyubiquitin ligase^80,81^ and protein kinase A^47^ in neurons, and mono-ubiquitination in β-cells^72^, has been shown to affect Tomosyn-1 function. Thus, we propose that Syt9 stabilizes Tomosyn-1 in the Syt9-Tomosyn-1-Stx1A non-fusogenic complex to prevent PM fusion of insulin granules. How nutritional and hormonal signaling pathways regulate Syt9 and Tomosyn-1inhibitory functions in affecting Stx1A-SNARE complexes formation remains part of future studies.

Increases in the formation of Stx1A-SNARE complexes and insulin secretion upon Syt9 knockdown could involve the role of distinct Syt isoforms. Multiple Syt isoforms can be present on the same granules, such as Syt1, Syt4, and Syt9. Also, Syt1 is present on granules that contain Syt7 and Syt9^82^. It is known that insulin granules harbor distinct isoforms of Syts, conferring different functional roles in exocytosis^82^. A recent study reported that Syt7 and Syt9 are present on distinct granules in β-cells^38^. Our data suggest that removing endogenous inhibitors Syt9 and Tomosyn-1 increases the availability of Stx1A to form SNARE complexes, facilitating insulin secretion (Figure 4). The formation of SNARE complexes requires the role of calcium sensor Syt isoforms that functions as a positive regulator of SNARE complex assembly, accelerating membrane fusion and insulin exocytosis. Thus, increases in the formation of Stx1A SNAREs upon Syt9 knockdown (Figure 4C) could involve the role of distinct Syt isoforms present on insulin granules with or without Syt9. Identifying Syt isoforms that facilitate the fusion of Syt9 granules and assessing if Syt9 insulin granules are (non)fusogenic requires further studies.

In conclusion, this work has identified Syt9 as an endogenous inhibitor of Stx1A-SNARE complexes to decrease insulin secretion. Insights into the mechanism demonstrate that Syt9 forms a non-fusogenic molecular complex with Tomosyn-1 and Stx1A, preventing insulin secretion. Further, the inhibitory effects of Syt9 on Stx1A-SNARE complex formation and insulin secretion are mediated by modulating the Tomosyn-1 protein abundance and binding ability to Stx1A.

## Materials and Methods

### Antibodies and chemicals

For the in-house insulin ELISA, the coating monoclonal anti-insulin antibody (clone D6C4) was from Fitzgerald (cat# 10R-I136A), insulin standard from EMD-Millipore (cat# 8013-K), detecting biotinylated anti-insulin antibody (clone D3E7) from Fitzgerald (cat#61R-I136bBT) and streptavidin-HRP from Pierce (cat#21126). Primary antibodies, β-actin (DSHB, cat#224-236-1), Stx1A (Sigma, cat#S0664), Syt5/9 (SYSY, cat#105053), Tomosyn-1 (SYSY, cat#183103), Vamp2 (SYSY, Cat#104008) and Snap25 (Biolegend, Cat#836304) were used in Western blotting, co-immunoprecipitation, and confocal imaging. The secondary antibodies used were goat antimouse (Cell Signaling, Cat#7076) and goat anti-rabbit (Jackson ImmunoResearch, Cat#111-035-003) for Western blotting and mouse Alexa Fluor 488 (Jackson ImmunoResearch, Cat#115-545-166) and rabbit Alexa Flour 647 (Jackson ImmunoResearch, Cat# 111-605-003) for confocal imaging.

### Expression constructs

The pcDNA3-m-Tomosyn-1construct was generously provided by Dr. Alexander Groffen, Virije Universiteit, Netherlands, and moloney murine leukemia virus-based retroviral vector (RVV, 3051) was a gift from Dr. Bill Sugden, University of Wisconsin, Madison. The m-Tomosyn-1 cDNA was subcloned to generate m-Tomosyn-1-V5 tagged RVV mammalian expression plasmid.

### Animals

The *Syt9*^-/-^ mice were generously given to us by Dr. Edwin Chapman, University of Wisconsin-Madison. Conditional *Syt9^Floxed^* mice obtained from Jackson Laboratory (Xu et al. 2007) were terminally crossed with E2a-CRE mice (Jackson Laboratory) to generate whole-body deletion of *Syt9*^-/-^ mice. All pups were weaned between 3-4 weeks of age. Male and female mice had free access to water, standard chow diet, and were housed in a temperature-controlled room with a 12 h light-dark cycle (6 AM-6 PM). All mice were kept following the University of Alabama at Birmingham Animal Research Program and NIH guidelines for the care and use of laboratory animals.

### Metabolic phenotyping

Body weight (BW), blood glucose, and plasma insulin were measured in *Syt9*^-/-^ and *Syt^+/+^* mice. Additionally, oral glucose tolerance test and *in vivo* insulin secretion were performed in response to oral glucose gavage (2 g glucose/kg BW of mice) after a 12 h fast. Tail vein blood was collected at different time points to determine blood glucose and plasma insulin levels. An insulin tolerance test was performed in response to an insulin dose (0.5U human insulin/BW of mice, Humulin R, Lilly, USA, Cat#002 8215-01) administered intraperitoneally to mice after a 6 h fast. Blood glucose levels were determined using the Contour Next blood glucose meter (Ascensia Diabetes Care, Switzerland). The plasma insulin concentration was measured by Ultra-Sensitive Mouse Insulin ELISA Kit (Crystal Chem, USA, Cat# 90080).

### Islet isolation and cell culture

Mouse islets were isolated from 8 to 10-week-old *Syt9*^-/-^ and *Syt9*^+/+^ mice using a collagenase digestion method and ficoll gradient procedure described previously^42,83^. Briefly, the pancreas was inflated by injecting collagenase (0.6 mg/ml) solution through the common bile duct, followed by removal, digestion steps, and ficoll gradient to obtain isolated islets. Handpicked islets were cultured overnight in supplemented RPMI 1640 culture medium containing 8 mM glucose for perifusion and static insulin secretion experiments.

INS1(832/13) β-cells, a gift from Dr. Christopher Newgard, Duke University, NC, were cultured in RPMI 1640 media supplemented with 10% heat-inactivated fetal bovine serum, 1 mM sodium pyruvate, 20 mM HEPES, 2 mM L-glutamine, and 100 units/ml of antimycotic-antibiotic along with 50 μM β-mercaptoethanol.

### Insulin secretion

Insulin secretion from mouse islets was performed as described previously^84 42^. Briefly, six size-matched islets from *Syt9*^+/+^ and *Syt9*^-/-^ mice were handpicked into each well of a 96-well plate. The insulin secretion assay was performed in a KRB-based buffer (Krebs-Henseleit Ringer bicarbonate buffer; KRB: 118.41 mM NaCl, 4.69 mM KCl, 1.18 mM MgSO4, 1.18 mM KH2PO4, 25 mM NaHCO3, 5 mM HEPES, 2.52 mM CaCl2, pH 7.4, and 0.5 % BSA). Islets were preincubated in 100 μl/well of KRB buffer containing 2.8 mM glucose. After 45 min, the preincubation buffer was replaced with the KRB incubation buffer containing insulin secretagogues. After 45 min, the incubation buffer was collected to estimate secreted insulin. Furthermore, islets were lysed in acid ethanol to determine cellular insulin content.

Insulin secretion from INS1(832/13) β-cells was performed as described previously^81^. Cells were seeded at a density of 100,000 cells/well in a 96-well plate at ~ 80% confluency. Forty-two-hours post-incubation, cells were washed and preincubated with 100 μl of the KRB-based buffer (118.41 mM NaCl, 4.69 mM KCl, 1.18 mM MgSO4, 1.18 mM KH2PO4, 25 mM NaHCO3, 20 mM HEPES, 2.52 mM CaCl2, pH 7.4, and 0.2% BSA) containing 1.5 mM glucose. After 2h, the preincubation buffer was replaced with the KRB incubation buffer containing insulin secretagogues. After 2h, an incubation buffer was collected to determine secreted insulin. Furthermore, cells were lysed in acid ethanol to determine cellular insulin content. Quantification of insulin was performed using in-house ELISA. The percent fractional insulin secretion was calculated as the amount of insulin secreted divided by the total insulin content.

### Islet perifusion

For the perifusion insulin secretion assay (as described^42^), a high capacity perifusion system from Bio Rep^®^ Perifusion was utilized. Approximately 75 islets were sandwiched in a chamber between two layers of Bio-Gel P-2 (Bio-Rad, Cat#1504118) bead solution (200 mg beads/ml in KRB buffer). Throughout the experiment, chambers containing islets and buffers were maintained at 37°C. Islets were perfused at a flow rate of 1 ml/min, and the flow-through containing secreted insulin was collected in a 96-well plate using an automatic fraction collector. The in-house ELISA was used to quantify cellular and secreted insulin.

### siRNA Transfection

Approximately 75–80% confluent INS1(832/13) cells were transfected with 0.2 μM of short interfering (si) RNAs, si-scramble (universal negative control, Sigma, Cat#SIC001, siSyt9 (sense-CAUUGUCCUGGAAGCUAAA, antisense-UUUAGCUUCCAGGACAAUG), si-Tomosyn-1 set 1(sense-GACCCAAAGCAGAAAGUUU, antisense-AAACUUUCUGCUUUGGGUC), and Tomosyn-1 set2 (sense-GAAAUUAAGCUUGCCAACU, antisense-AGUUGGCAAG CUUAAU UUC) using Lipofectamine 2000 (Invitrogen) at 1:2 ratio in Opti-MEM. After 6 h, transfection solution was replaced with RPMI 1640 supplemented medium without antibiotics. After 42 h, cells were harvested for insulin secretion, Western blotting, or qPCR.

### Subcellular fractionation

INS1(832/13) cells were washed with ice-cold 1X PBS and harvested in HEPES/EGTA buffer (20 mM HEPES,1 mM EGTA, 1 mM PMSF, and protease inhibitor cocktail tablet) using a cell scraper. The cell lysate was homogenized, passing through a 22G needle 15-20 times to break up the cell membrane while keeping the granule and nuclear membrane intact. The homogenized suspension was centrifuged at 1000xg for 5 min at 4°C to remove nuclear protein and debris. The supernatant was collected and centrifuged again to make sure all heavy contaminants were removed. Then supernatants were carefully placed on a 10 ml sucrose gradient (top to bottom layers formed with the concentration of sucrose of 0.45, 0.60, 0.75, 0.90,0.1.05,1.20,1.35,1.50, 1.65, 1.80 and 2.00) in Beckman high speed centrifuge tubes for rotor SW41 Ti. Ultracentrifugation was performed at 111,000xg for 18 hours at 4°C. Fractions were collected from the top, and an equal amount of protein from each fraction was separated on SDS-PAGE, transferred to PVDF membranes, immunoblotted with their respective antibodies, and developed with ECL.

### Immunoprecipitation

INS1 (832/13) cells were seeded in a 100 mm tissue culture dish at 60% confluency. Transient transfection was performed using 0.2 μM siRNA or 10 μg Tomosyn-1-V5-RVV or GFP-V5-RVV using Lipofectamine 2000 (Invitrogen) construct at a 1:2 ratio in Opti-MEM. Forty-eight-hour post-transfection, cells were pre-incubated for 2 h in KRB containing 1.5 mM glucose before 40 mM KCl treatment for 15 min.

#### Immunoprecipitation of overexpressed Tomosyn-1

Cell lysates were prepared using 1ml IP lysis buffer (25 mM Tris-HCl, pH 7.5, 50 mM NaCl, 2mM MgCl2, 1 mm Na2EDTA, 1 mM EGTA, 1mM CaCl2, 0.5% Triton X-100, 2.5 mM sodium pyrophosphate, 1 mM β-glycerophosphate, 1 mM Na3VO4, 1 mM PMSF and protease inhibitor cocktail tablet). Protein concentration was estimated using BCA (Pierce BCA protein assay kit, Cat#PI23227). Immunoprecipitation was performed in IP lysis buffer supplemented with 0.2% BSA using 50 μl of equilibrated magnetic bead suspension (anti-V5-tag mAb-Magnetic Beads; MBL life sciences M167-11). The immunoprecipitated complex was washed thrice with 1ml IP lysis buffer with 0.2% BSA. The protein complex was eluted in 80 ml 2.5X Laemmli buffer with 1 mM DTT followed by heating at 95°C for 5 min. Proteins were separated by 10 % SDS-PAGE gel.

#### Immunoprecipitation of endogenous proteins

Cells combined from two 10 cm dishes were lysed in 700 μl IP lysis buffer (25 mM Tris-HCl, pH 7.5, 50 mM NaCl, 2mM MgCl2, 1 mM Na2EDTA, 1 mM EGTA, 1mM CaCl2, 0.5% NP-40, 0.1% SDS, 2.5 mM sodium pyrophosphate, 1 mM Na3VO4, 1 mM PMSF, and protease inhibitor cocktail tablet) by placing on a rotating shaker for 1 hour at 4°C followed by a centrifugation at 13000 rpm for 10 min. The precleared lysate was incubated overnight with Stx1A antibody (2 mg/ml) or IgG isotype control. The next day, 50 ml of equilibrated magnetic beads were added to the reaction mixture and incubated at 4°C for 1 h. The immunoprecipitated complex was washed three times with 1 ml IP lysis buffer and eluted in 80 ml 2.5X Laemmli buffer with1 mM DTT, followed by heating at 95°C for 5 min. Proteins were separated by 10 % and 12% SDS-PAGE gels and transferred to a PVDF membrane followed by blotting with Clarity Western ECL substrate (Bio-Rad, cat#1705061).

### Western blot

Approximately 100 mg of brain tissue from 6-8-week-old *Syt9* and *Syt9*^-/-^ mice were pulverized using a mortal pestle on dry ice and added to ice-cold 1 ml lysis buffer (20 mM Tris-HCl pH 7.5, 150 mM NaCl, 1 mM Na2EDTA, 1 mM EGTA, 1% Triton X-100, 2.5 mM sodium pyrophosphate, 1 mM β-glyceraldehyde, 1 mM Na3VO4, 1 mM NaF, 1 mM PMSF, and protease inhibitor cocktail). Samples were homogenized, sonicated, subjected to high-speed spin, and quantified by BCA assay for Western blotting (described above). Similarly, islets (400 per biological replicate) isolated from 6-8-week-old *Syt9*^+/+^ and *Syt9*^-/-^ mice were added to the 200 μl lysis buffer. Cell lysates were sonicated and subjected to high-speed spin, and BCA quantified soluble fractions were used for Western blotting. Similarly, INS1 lysates were prepared. The following primary antibodies were used: anti-actin (1:5,000), anti-Stx1A (1:10,000), anti-Syt5/9 (1: 3,000), and anti-Tomosyn-1 (1:5,000). Secondary antibodies used were goat anti-mouse (1:10,000) and donkey anti-rabbit (1:10,00).

### Immunohistochemical analysis

INS1 (832/13) cells cultured on poly-D-lysine (50 mg/ml) coated coverslips were preincubated for 2 h in KRB buffer containing 1.5mM glucose before stimulation with 40 mM KCl for 15 min. Cells were fixed with 4% paraformaldehyde (2ml/well) in 37°C for 20 minutes, washed five times at 5 min intervals with 1X PBS, blocked with 10% normal donkey serum for 1 h, and permeabilized (1X PBS, 2.5% normal goat serum, 2.5% normal donkey serum, 1% BSA, 0.15% Triton X-100) for 30 minutes. After washing (1X PBS, 3 times at 5 min intervals), primary antibody labeling was performed for 3 hours by placing the coverslips in a humidity chamber. After 1XPBS washes, secondary antibody labeling was performed in the dark for one hour. Finally, the coverslip was mounted on a microscope slide with a mounting buffer containing DAPI. Confocal imaging was performed using LSM750 Zeiss at 60X magnification. Images were analyzed with ImageJ, and colocalized pixels were plotted and expressed as Manders’ and Spearman’s coefficient values. The primary antibodies used were anti-Stx1A (1:500) and anti-Tomosyn-1 (1:500) and anti-Syt5/9 (1: 500). The secondary antibodies were mouse Alexa fluor 488 (1:1,000) and rabbit Alexa fluor 647 (1:1,000).

### Quantitative real-time PCR

Total RNA was harvested from mouse islets and INS1(832/13) cells by using a QIAGEN RNeasy Plus Kit (Cat# 74034) and a TRIzol™ reagent (Invitrogen, cat#15596018) respectively. Following RNA extraction, cDNA was synthesized using a high-capacity cDNA reverse transcription Kit (Applied Biosystems, cat# 4368814). The mRNA abundance for the genes (Supplementary Table 1) of interest was determined by quantitative PCR using Fast Start SYBR Green (Roche, Cat# 4673484001). The data was quantified by the comparative ΔCT method and expressed relative to β-actin mRNA.

### Statistical Analysis

Data are represented as means ± SEM. Statistical significance was performed using Student’s twotailed unpaired t-test for independent data. The significance limit was set at p < 0.05.

## Author contributions

M.M.R., S.B., writing and editing of the manuscript; M.M.R., S.B., A.P., A.M., J.A., for reviewing of the manuscript; M.M.R., S.B., A.P., for data curation; M.M.R., A.P., K.S., SB., investigations; M.M.R., S.B., A.P., analyzed and data interpretation; H.A., S.B., J.E.K., E.C., provided resources for the project; S.B., conceptualization of the project; S.B., supervision of the project; S.B., funding for the project; S.B., project administration. All authors provided critical feedback and helped shape the research, analysis, and manuscript.

## Acknowledgments

We thank Drs. Sasanka Ramanadham and Jeonga-a Kim for critical feedback on the project and manuscript.

## Funding

This work was also supported by National Institutes of Health NIDDK Grants 4 R00 DK95975-03, R01DK120684, 1R21DK129968-01, and Diabetes Research Center (DRC) Grant P30DK079626-10 (to S.B). *The authors declare that they have no conflicts of interest with the contents of this article*. The content is solely the responsibility of the authors and does not necessarily represent the official views of the National Institutes of Health.

## Notes

### Competing Interest Statement

The authors have declared no competing interest.

