## Supplemental Data for "Synaptotagmin-9 and Tomosyn-1 molecular complex regulates Stx1A SNAREs to inhibit insulin secretion from pancreatic β-cells"

Affiliations:

**Summary**

Stimulus-coupled insulin secretion from b-cells involves the fusion of insulin granules to the plasma membrane (PM) via SNARE complex formation—a cellular process key for maintaining whole-body glucose homeostasis. Optimal insulin secretion depends on how the clamping of SNAREs is released, rendering granules fusogenic. We show that an insulin granule protein synaptotagmin-9 (Syt9) deletion in lean mice increased glucose clearance, random-fed plasma insulin levels, and insulin secretion (in vivo and ex vivo islets) without affecting insulin sensitivity. These outcomes demonstrate that Syt9 has an inhibitory function in insulin secretion. Moreover, Syt9 interacts with PM-Stx1A and soluble Tomosyn-1 proteins to form non-fusogenic complexes between PM and insulin granules, preventing Stx1A-SNARE formation and insulin secretion. Furthermore, Syt9 inhibits SNARE-complex formation by posttranscriptional regulation of Tomosyn-1. We conclude that Syt9 and Tomosyn-1 are endogenous inhibitors that modulate Stx1A availability to determine b-cell secretory capacity.

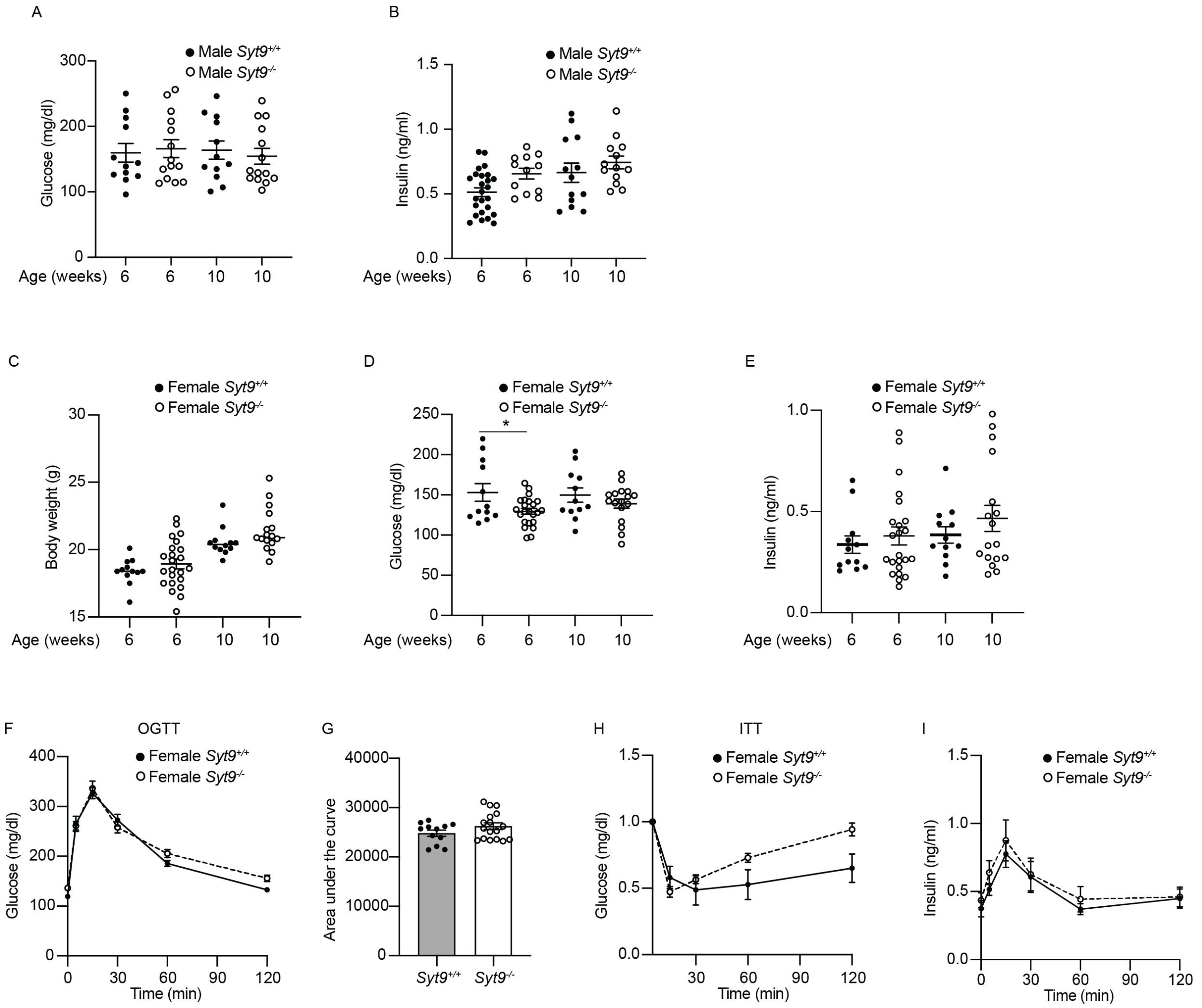

**Figure S1:** Validation of whole body *Syt9* $^{-/-}$ mice and metabolic analyses in both male and female mice.

(A) Plasma glucose levels (mg/dl) were determined after fasting in male mice (n > 10).

(B) Plasma insulin levels (ng/ml) were determined after fasting in male mice (n > 10).

(C) Body weight determined for *Syt9*$^{+/+}$ and *Syt9*$^{-/-}$ mice at 6 and 10 weeks of age (n > 10).

(D) Plasma glucose levels (mg/dl) were determined after fasting in female mice (n > 10).

(E) Plasma insulin levels (ng/ml) were determined after fasting in female mice (n > 10).

(F-G) Oral glucose tolerance test (OGTT) and corresponding area under the curve in *Syt9*$^{+/+}$ and *Syt9*$^{-/-}$ female mice (n > 10).

(H) Insulin tolerance test (ITT) was performed in *Syt9*$^{+/+}$ and *Syt9*$^{-/-}$ female mice (n > 5).

(I) In vivo insulin secretion during OGTT in *Syt9*$^{+/+}$ and *Syt9*$^{-/-}$ female mice (n > 10).

Data are presented as mean ± SEMs. *P < 0.05; **P < 0.01; ***P < 0.001

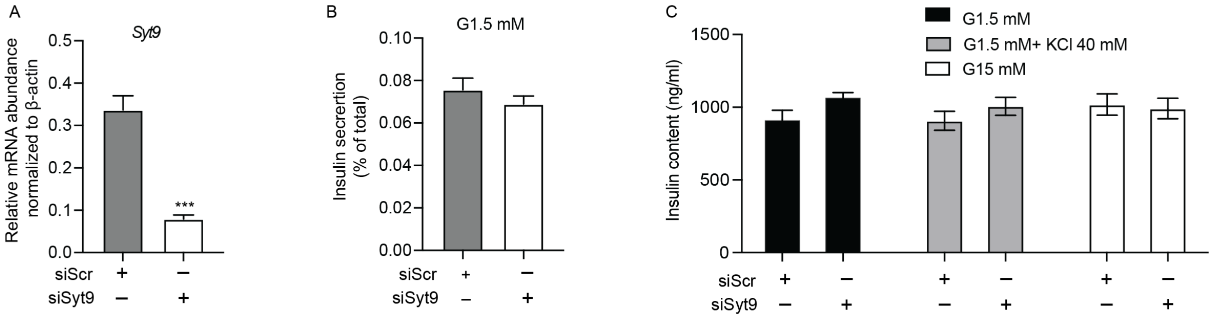

**Figure S2:** The siRNA-mediated knockdown of Syt9 in INS1(832/13) cells has no effect on basal insulin secretion and total insulin content.

(A) Relative mRNA expression of Syt9 in INS1(832/13) cells transfected with siScramble (siScr) or siSyt9 (n = 4).

(B) Basal insulin secretion in INS1 (832/13) cells transfected with siScramble (siScr) or siSyt9 (n = 3).

(C) Insulin content in response to low glucose (1.5 mM), high glucose (15.0 mM) and KCl (40 mM at 1.5 mM glucose) in INS1(832/13) cells transfected with siScr or siSyt9 (n = 3).

Data are presented as mean ± SEMs. $*P < 0.05$; $**P < 0.01$; $***P < 0.001$

Figure S3

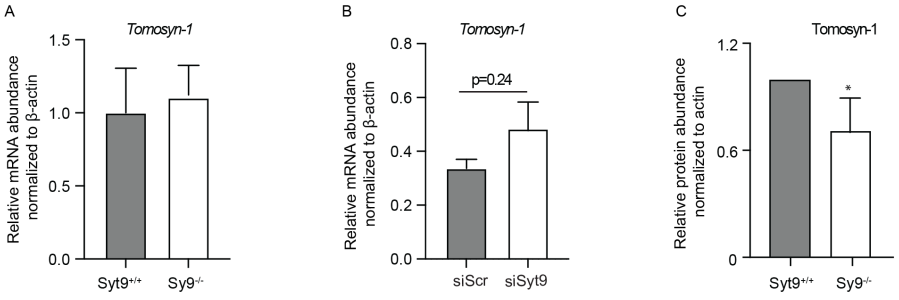

**Figure S3:** The effect of Syt9 loss in Tomosyn-1 mRNA and/or protein level

(A) Relative mRNA expression of Tomosyn-1 in *Syt9*$^{+/+}$ and *Syt9*$^{-/-}$ mouse islets (n = 4).

(B) Relative mRNA expression of Tomosyn-1 in INS1(832/13) cells transfected with siScr or siSyt9 (n = 4).

(C) Tomosyn-1 protein abundance in whole brain of *Syt9*$^{+/+}$ vs. *Syt9*$^{-/-}$ mice (n=4).

Data are presented as mean ± SEMs. $^{*}P < 0.05$; $^{**}P < 0.01$; $^{***}P < 0.001$

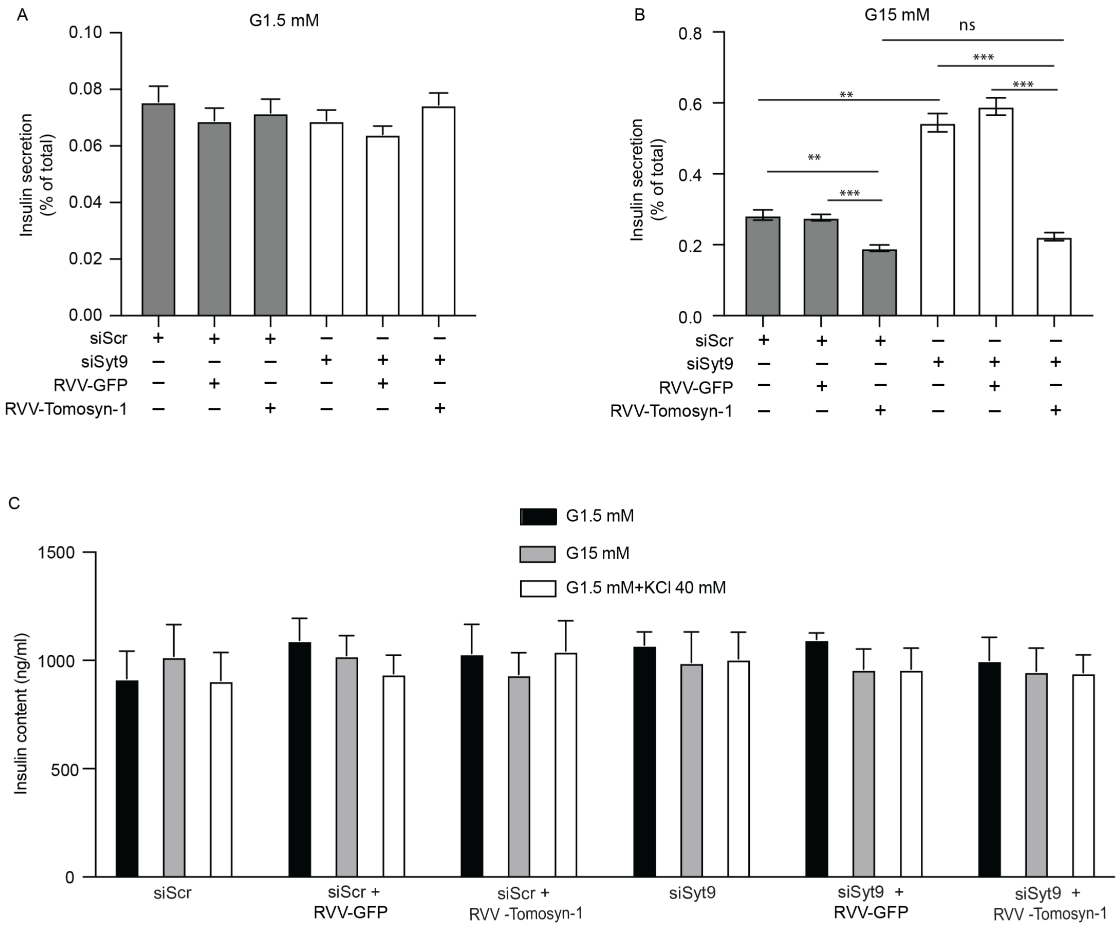

**Figure S4:** Insulin secretion in response to different stimuli in cells having either overexpressed Tomosyn-1 with or without knockdown of Syt9.

(A) Insulin secretion in response to basal glucose (1.5 mM) from INS1(832/13) cells (n = 3).

(B) Insulin secretion in response to high glucose (15 mM) from INS1(832/13) cells (n = 3).

(C) Insulin content in response to basal, high glucose, and KCl in INS1(832/13) cells (n = 3).

Data are presented as mean ± SEMs. *P < 0.05; **P < 0.01; ***P < 0.00

**Table 1: The list of key reagents used in this study**

| Name | Source | Catalog No. | Description |
|---|---|---|---|
| **Antibody** | | | **Dilution** |
| Rabbit- anti-Synaptotagmin-9 | Synaptic System | 105 053 | 1:3000 |
| Rabbit-anti-Tomosyn-1 | Synaptic System | 183103 | |
| Mouse-anti-Syntaxin 1 | Sigma | S0664 | 1:10,000 |
| Mouse-anti-SNAP25 | Biolegend | 836304 | 1:3000 |
| Rabbit- anti-Vamp2 | Synaptic System | 104 008 | 1:3000 |
| Mouse-anti-β- actin | DSHB | 224-236-1 | 1:10,000 |
| Peroxidase Goat Anti-Rabbit IgG (H+L) | Jackson Immune Research | 111-035-003 | 1:10,000 |
| Mouse Alexa Fluor 488 | Jackson ImmunoResearch | 115-545-166 | 1:1000 |
| Rabbit Alexa Flour 647 | Jackson ImmunoResearch | 111-605-003 | 1:1000 |
| **Reagents** | | | |
| Ultra-Sensitive Mouse Insulin ELISA Kit | Crystal Chem, USA | 90080 | |
| Humulin R Regular Insulin U | Lilly, USA | 002 8215-01 | |
| Bio-Gel P-2 | Bio-Rad | 1504118 | |
| TPA | | | |
| MISSION® siRNA Universal Negative Control | Sigma | SIC001 | |

| **siRNA** | Sense sequence (5'-3') | Antisense sequence (5'-3') |
|---|---|---|
| Syt9 | CAUUGUCCUGGAAGCUAAA | UUUAGCUUCCAGGACAAUG |
| Tomosyn-1_S1 | GACCCAAAGCAGAAAGUUU | AAACUUUCUGCUUUGGGUC |
| Tomosyn-1_S2 | GAAAUUAAGCUUGCCAACU | AGUUGGCAAGCUUAAUUUC |
| **qPCR Primers** | | |
| Beta-actin | TGTGATGGTGGGAATGGGTCAGAA | TGTGTTGCCAGATCTTCTCCA TGT |
| Syt-9 | CAGACCCTGAATCCACACTTT | GGAGAAGCGATCGAAGTCATA C |
| Tomosyn-1 | TGCAAGACTGTTCGCCATGGA T | GCTGTCGTGCTGGCAATAACAT |
